# The Oral Bioavailability of Lurasidone is Impacted by Changes to the Gut Microbiome: Implications for Antipsychotic Therapy

**DOI:** 10.1101/2024.07.17.604016

**Authors:** Kate Collins, Srinivas Kamath, Tahlia R. Meola, Anthony Wignall, Paul Joyce

## Abstract

Atypical antipsychotics are crucial for the management of schizophrenia and bipolar disorder, yet they exhibit significant pharmacokinetic variability which leads to inconsistent therapeutic responses. This study investigates the hypothesis that gut microbiome composition critically influences the oral bioavailability of lurasidone, a poorly soluble weak base antipsychotic with pH-dependent solubility. To investigate this, male Sprague-Dawley rats underwent systematic gut microbiome manipulation through pretreatment with antibiotics or prebiotics (inulin) for 14 days prior to a single oral dose of lurasidone. Pharmacokinetic analysis of collected plasma samples revealed a significant 4.3-fold increase in lurasidone bioavailability following prebiotic pretreatment, compared to a control (no pretreatment) group. Conversely, lurasidone bioavailability was highly variable in rats with a depleted microbiome (*i.e.*, antibiotic treatment group), with 80% of animals demonstrating lower bioavailability than the control group. Characterisation of gut microbiome composition and short-chain fatty acid (SCFA) concentrations demonstrated positive correlations between lurasidone bioavailability, microbial diversity, and SCFA levels, mediated by modulation of luminal pH. Elevated SCFA levels created a favourable environment for lurasidone solubilisation by lowering intestinal pH. These findings highlight the potential for optimising antipsychotic pharmacokinetics through personalised microbiome interventions. Furthermore, the correlation between SCFAs and lurasidone bioavailability suggests their potential as biomarkers for predicting inter-patient pharmacokinetic variability, particularly for poorly soluble weak bases. Thus, new avenues are opened for developing novel co-therapies and screening tools to enhance antipsychotic pharmacokinetic performance, potentially improving treatment outcomes for patients with schizophrenia and bipolar disorder.

**Graphical Abstract:** 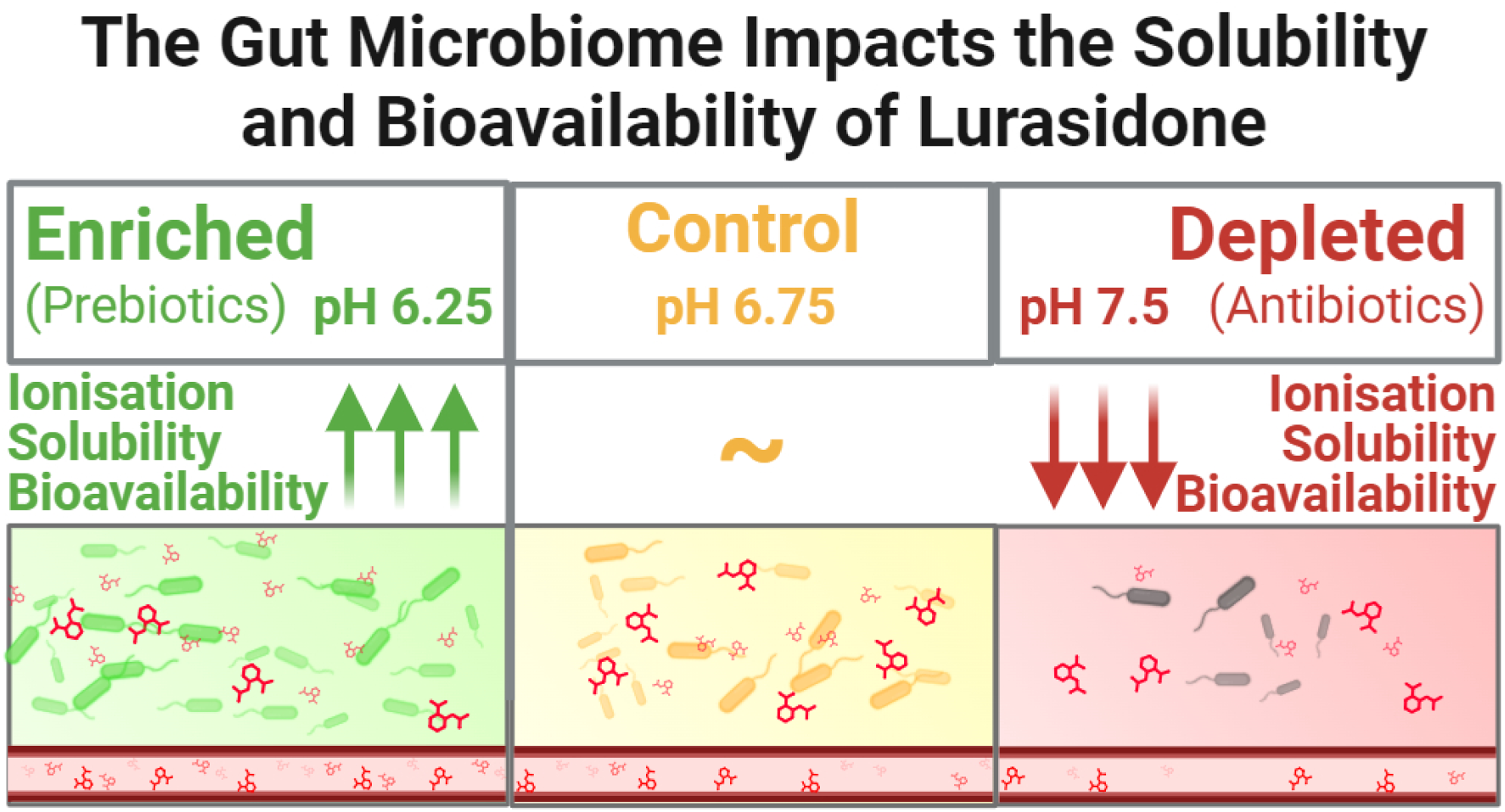

## 1. Introduction

Atypical antipsychotics are vital pharmacotherapeutics for a broad spectrum of mental health conditions, including schizophrenia, bipolar disorder, major depressive disorder and anxiety [1–3]. However, their effectiveness is often compromised by severe side effects and variable pharmacokinetics, leading to poor patient outcomes, low response rates, and treatment abandonment. Despite this, antipsychotics continue to be prescribed without full consideration of their potential negative effects on metabolic function and overall mental health [1,4–6].

A critical, yet frequently overlooked aspect of antipsychotic treatment is the bidirectional interaction between these drugs and the gut microbiome [1,3,7,8]. This bidirectional relationship manifests in two vital ways:

1) Antipsychotics and their excipients can disrupt the gut microbiome, leading to adverse metabolic effects such as weight gain, hyperglycaemia, and hyperlipidaemia [4,5,7,9,10].
2) The gut microbiome can alter the pharmacokinetics of orally administered drugs, impacting their absorption, biodistribution, metabolism, and excretion [1,3,8,9].

These bidirectional interactions ultimately contribute to the side effect profile experienced by many patients and form the underlying mechanisms behind the significant interindividual variability observed in antipsychotic treatment responses [1,3,8,11,12].

Lurasidone, a benzothiazole derivative atypical antipsychotic and poorly soluble weak base (PSWB) (pKa = 7.6), serves as a compelling example of these drug-gut interactions due to its pH-dependent solubility [9,10,13,14], poor and variable oral bioavailability (in the reported range of 9-19%), and significant “food effect” where bioavailability varies up to 3.5-fold depending on the consumption of food [9,15,16]. Prior literature by Cussotto et al. demonstrated that microbiome depletion in adult male Sprague-Dawley rats (n: 6-7/group, ampicillin 1g/L, vancomycin 500mg/L, and imipenem 250mg/L for 14 days in drinking water) led to an 82% increase in olanzapine bioavailability, while the pharmacokinetics of risperidone was unaltered by antibiotic pretreatment [6]. The cause for microbiome-mediated changes in olanzapine pharmacokinetics were not well-understood, but a negative correlation was shown between olanzapine bioavailability and the abundance of *Alistipes*, a butyrate-producing Gram-negative genus found in the gut microbiome. Ultimately, this discrepancy underscores the complexity and drug-specific nature of these interactions, which may be influenced by factors such as enzymatic degradation, and drug metabolism, as summarised by Minichino et al. in a psychotropic context [3].

Based on previous findings in literature, a hypothesised mediator in this intricate relationship is the production of short-chain fatty acids (SCFAs) by the gut microbiome. These metabolites, primarily acetate, propionate, and butyrate, are produced through microbial fermentation of indigestible dietary fibres and have been shown to influence the pH of the intestinal lumen [17,18]. While several studies have demonstrated that enrichment of the gut microbiome enhances SCFA production [19–22], the specific implications for individual drugs remain poorly understood and explored. For PSWBs with pH-dependent solubilities, like lurasidone, it is expected that microbiome- and SCFA-mediated changes to luminal pH will lead to significant pharmacokinetic variability due to changes in the relative degree of drug solubilisation within the small intestine.

This novel study aims to address this critical gap in our understanding of antipsychotic pharmacokinetics by systematically investigating the impact of the gut microbiome on lurasidone pharmacokinetics. To achieve this, Sprague-Dawley rats received one of three pretreatments prior to lurasidone administration: 1) PBS (control), 2) antibiotics (ampicillin, vancomycin, and imipenem), or 3) prebiotics (inulin). Pharmacokinetic analyses were undertaken in combination with microbiota and metabolome profiling with the objective of unravelling the variable and poor oral bioavailability of lurasidone. Ultimately, the insights gained from this study may pave the way for more personalised and effective treatment strategies that mitigate pharmacokinetic variability by maximising oral bioavailability with the intention of improving therapeutic outcomes for patients with mental health disorders [12]. Furthermore, these findings may have far-reaching implications for optimal drug usage across various classes of psychotropic medications, ultimately contributing to more informed and nuanced approaches in psychopharmacology. These findings may lead to the development of predictive pharmacokinetic models based on gut microbiome composition, advancing the field towards truly personalised mental health treatment.

## 2. Materials and methods

### 2.1 Animal Ethics & Husbandry

The *in vivo* study received approval from the Animal Ethics Committee at the University of South Australia (approval #U24-21), adhering to the NIH’s Principles of Laboratory Animal Care (NIH publication #85-23), revised in 1985) and the Australian code for the care and use of animals for scientific purposes (8^th^ edition 2013, revised 2021), and was conducted and reported following ARRIVE guidelines. The study was conducted on 8-week-old male Sprague-Dawley rats sourced from Ozgene (Canning Vale, Australia) with a mean starting weight of 270-330 g. Rats were housed in groups of 2 in a temperature-, humidity-, pressure-controlled animal facility with a 12h/12h light/dark cycle with access to standard rodent chow and water. Only healthy rats were included in the study, and those showing signs of illness or abnormal behaviour during the acclimation period were excluded. Randomisation and blinding were implemented during the allocation of animals to treatment groups and throughout the data collection process. The researchers administering treatments, as well as those assessing outcomes, were blinded to the group allocations to prevent bias.

### 2.2 In vivo study design

The oral bioavailability of lurasidone (Hangzhou Dayangchem Co. Ltd., Hangzhou City, China) was investigated in three randomly assigned treatment groups (*n* = 5). The sample size for each treatment group (*n* = 5) was based on Power calculations for anticipated changes in the lurasidone area-under-the-curve (AUC), using a power level of ≥ 0.8 and a significance level of 0.05. A single-gender was used for the pharmacokinetics study to minimise gender-based variation commonly observed in oral pharmacokinetics investigations. Treatment groups were as follows: 1) control group where rats were provided *ad libitum* access to standard drinking water; 2) enriched microbiome group where rats were provided *ad libitum* access to drinking water with inulin dissolved (100 mg/L; average degree of polymerisation of 35 purchased from Merck, Australia); and, 3) depleted microbiome group where rats were provided *ad libitum* access to drinking water with vancomycin (500 mg/L), ampicillin (1 g/L), and imipenem (250 mg/L) dissolved (antibiotics purchased from Hangzhou Dayangchem Co. Ltd., Hangzhou City, China). Following 14 days of pre-treatment with prebiotics (*i.e.,* inulin) or antibiotics to manipulate the gut microbiome, each treatment group was administered a single dose of lurasidone (20 mg/mL in PBS) via oral gavage. Blood samples were collected over various time points, (0, 0.33, 0.66, 1, 1.5, 2, 4, 6, 8, 24h) for pharmacokinetic analysis. After the final samples were collected, rats were anesthetised using 5% isoflurane, followed by cardiac puncture and cervical dislocation.

### 2.3 Pharmacokinetic analysis

Blood samples were collected from the saphenous vein and centrifuged to collect the plasma. For LCMS analysis the plasma was diluted with acetonitrile (5:9 ratio) and spiked using an internal standard (ziprasidone; 10μl). Pharmacokinetic analyses were performed using Phoenix® WinNonlin® Version 8.3 (Pharsight®, a Certara™ company), using non-compartmental methods (Model 200–202, Extravascular Input) and a linear-trapezoidal approach.

### 2.4 Gut microbiota analysis

Following treatment the study completion, animal caecal contents were collected and stored in a -80°C freezer for future analysis. Samples were sent to the Australian Genomics Research Facility (Brisbane, Australia) for DNA extraction, PCR amplification, and 16S rRNA sequencing. Using the Quantitative Insights to Microbiology Ecology (QIIME 1.8and the Silvia reference database the 16S rRNA sequences were processed for the V3-V4 hypervariable regions, with raw reads clustered into operational taxonomic units (OTUs). OTUs were assigned taxonomy using the QIAGEN Microbial Insights – Prokaryotic Taxonomy Database (QMI-PTDB) within the QIAGEN CLC Genomics Workbench version 23.0.4 (Hilden, Germany). Alpha diversity was measured via Shannon’s Index at the genus level, and beta diversity analyses were conducted using Principal Coordinate Analyses (PCoAs) based on Bray-Curtis’ dissimilarity metrics. Statistical significance was evaluated through Permutational Multivariate ANOVA (PERMANOVA) on the beta diversity plots.

### 2.5 Short-chain fatty acid analysis

SCFA concentration in caecal contents was analysed using a modified gas chromatography-mass spectrometry (GCMS) protocol (Moreau et al, 2003), modified for the GC–MS (7890A GC System and 5975C inert XI EI/CI MSD with an EI inert 350 source) and MassHunter Workstation Software (MassHunter, Agilent Technologies). The protocol details have been previously published [23].

### 2.6 Intestinal pH analysis

Following the study completion, small intestinal media was collected from each rat for quantifying intestinal pH. Due to the relatively low volumes of small intestinal media, the media comprises that collectively obtained from the duodenum, jejunum, and ileum. The pH of the small intestinal media was quantified at 37°C using a HALO® wireless pH meter with microbulb (HI10832, Hanna Instruments, Keysborough, Australia).

### 2.7 In vitro lurasidone drug solubility

Lurasidone drug solubilisation was analysed at varying pH levels within simulated fasted state intestinal fluid (FaSSIF; Biorelevant.com Ltd, London, UK) to predict the impact of microbiome-mediated changes in intestinal pH on drug solubilisation. The pH of FaSSIF was adjusted to 6.25, 6.75, and 7.5 using 0.1M NaOH or HCl. Lurasidone was added to each FaSSIF media in excess (*i.e.*, 10 mg/mL) and gently stirred overnight at 37°C. Samples were centrifuged at 20,000 *g* for 20 minutes at room temperature to separate precipitated/insoluble lurasidone. The supernatant was collected and analysed using liquid chromatography-mass spectrometry (LCMS) to quantify the soluble fraction within each FaSSIF media. The protocol details have been previously published [10]. Solubility studies were performed in triplicate.

### 2.8 Statistical Analysis

GraphPad Prism Version 8.0 (GraphPad Software Inc., California) was used for all statistical analyses (excluding 16S sequencing analyses) of experimental data. An unpaired t-test and one-way ANOVA were used to determine statistically significant differences, followed by Tukey’s post-test for multiple comparisons. Values are expressed as the mean ± standard deviation (SD), with statistical significance set at p < 0.05. In the figures, statistical significance is denoted by * p < 0.05, ** p < 0.005, *** p < 0.0005, and **** p < 0.00005.

## 3. Results

### 3.1 Prebiotics and antibiotics differentially impact gut microbiota diversity

An overview of the dosing regimen and study design to investigate the impact of the gut microbiome on lurasidone bioavailability is presented in **Fig. 1A**. The 14-day pre-treatment approach comprised of treatment groups being administered either 1) no pre-treatment (*i.e.*, control), 2) antibiotics, or 3) prebiotics *ad libitum* in drinking water. Analysis of microbial abundance at the family taxonomic level from the caecal contents from each animal, illustrated in **Fig. 1B**, with grouped data highlighted in **Fig. 1C**, indicates that the pre-treatment corresponds to treatment groups with a 1) control microbiota, 2) depleted microbiota, and 3) enriched microbiota. Microbial abundance increased following prebiotic pre-treatment, in particular members of the *Lachnospiraceae* and *Bacteroidacae* families. In contrast, *Enterobacteriaceae* show predominance in the antibiotic-treated group. Observed changes in total operation taxonomic units (OTUs) and alpha diversity (measured by Shannon’s index) are presented in **Fig. 1D** and **Fig. 1E** for the three groups, where the prebiotic treatment group exhibited an enriched microbiota through increases in OTUs (+50 mean increase in OTUs) and Shannon’s index (+1 mean increase in Shannon’s index) compared to the control group. Contrary to this, the antibiotic treatment group displays a significant decrease in OTUs (-50 OTUs) and Shannon’s index (-1 decrease), when compared to the control group. PERMANOVA of principal coordinate analysis (PCoA) reveals statistically significant differences in beta diversity between the three treatment groups (Fig 1F), further highlighting that three distinct microbial populations were present for the respective treatment groups.

**Figure 1.**
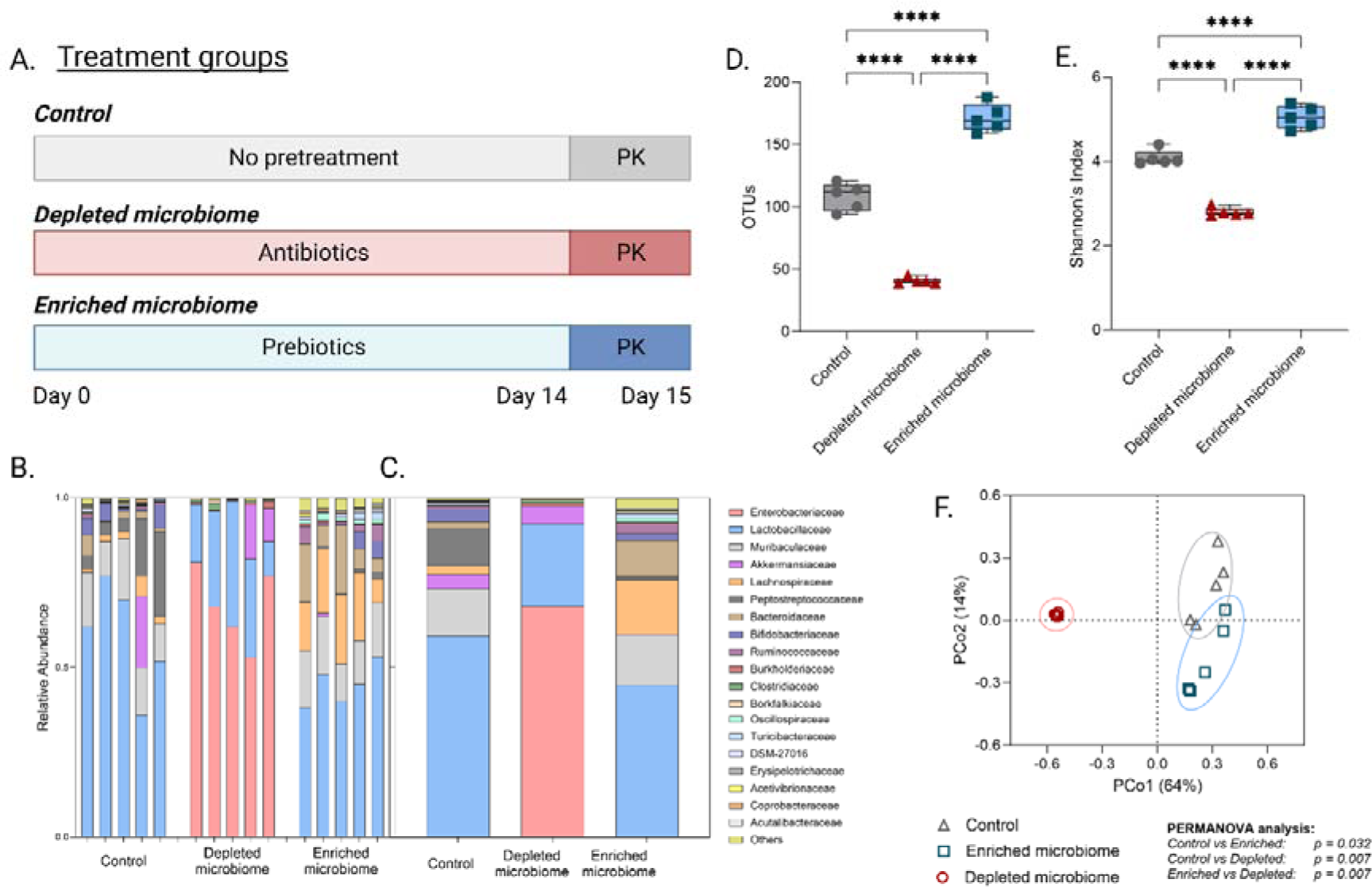
Microbial pretreatment resulted in three distinct microbiota compositions between the treatment groups. [**A**] A schematic representation of the treatment protocols for each group. Relative abundance of gut microbiota families post-treatment for individual animals (Panel [**B**]) and grouped averages (Panel [**C]**). Changes in OTUs (Panel [**D**]) and Shannon’s Index (Panel [**E**]) reveal statistically significant changes in microbial diversity across the three treatment groups. Principal coordinate analysis demonstrates statistically significant shifts in microbiota composition across the three treatment groups (Panel [**F**]). Statistical significance was assessed using unpaired t-test and one-way ANOVA for α-diversity and PERMANOVA for β-diversity. The asterisk (****) denotes statistically significant results (p < 0.0001).

### 3.2 Microbiota diversity positively correlates with caecal SCFA levels

The concentration of SCFAs, specifically acetic, propionic, and butyric acid, was quantified within caecal contents to determine the impact of microbiota manipulation on the metabolome. Prebiotic and antibiotic pre-treatment resulted in a significant difference (*p* ≤ 0.001) across all SCFA levels, including the total SCFA concentration (**Fig. 2A-D**). The increase in each individual SCFA observed for the enriched microbiome group (*i.e.*, prebiotic treatment) was increased, yet not statistically significant, compared to the control group. In contrast, a statistically significant depletion was observed for the antibiotic treatment group across each SCFA. Prebiotic pre-treatment leading to an enriched microbiome revealed the greatest total SCFA concentration, between 200-300μmol g^-1^, while antibiotic pretreatment leading to a depleted microbiome revealed the lowest total SCFA concentration (< 50 μmol g^-1^). A strong positive correlation between total SCFA (μmol g^-1^) and Shannon’s index (R^2^ = 0.93) is displayed in **Fig. 2E**, indicating that microbial diversity leads to an increase in total SCFA concentration. Similarly, a strong positive correlation (R^2^ = 0.83) can be noted between total SCFA (μmol g^-1^) and OTUs (**Fig. 2F**).

**Figure 2.**
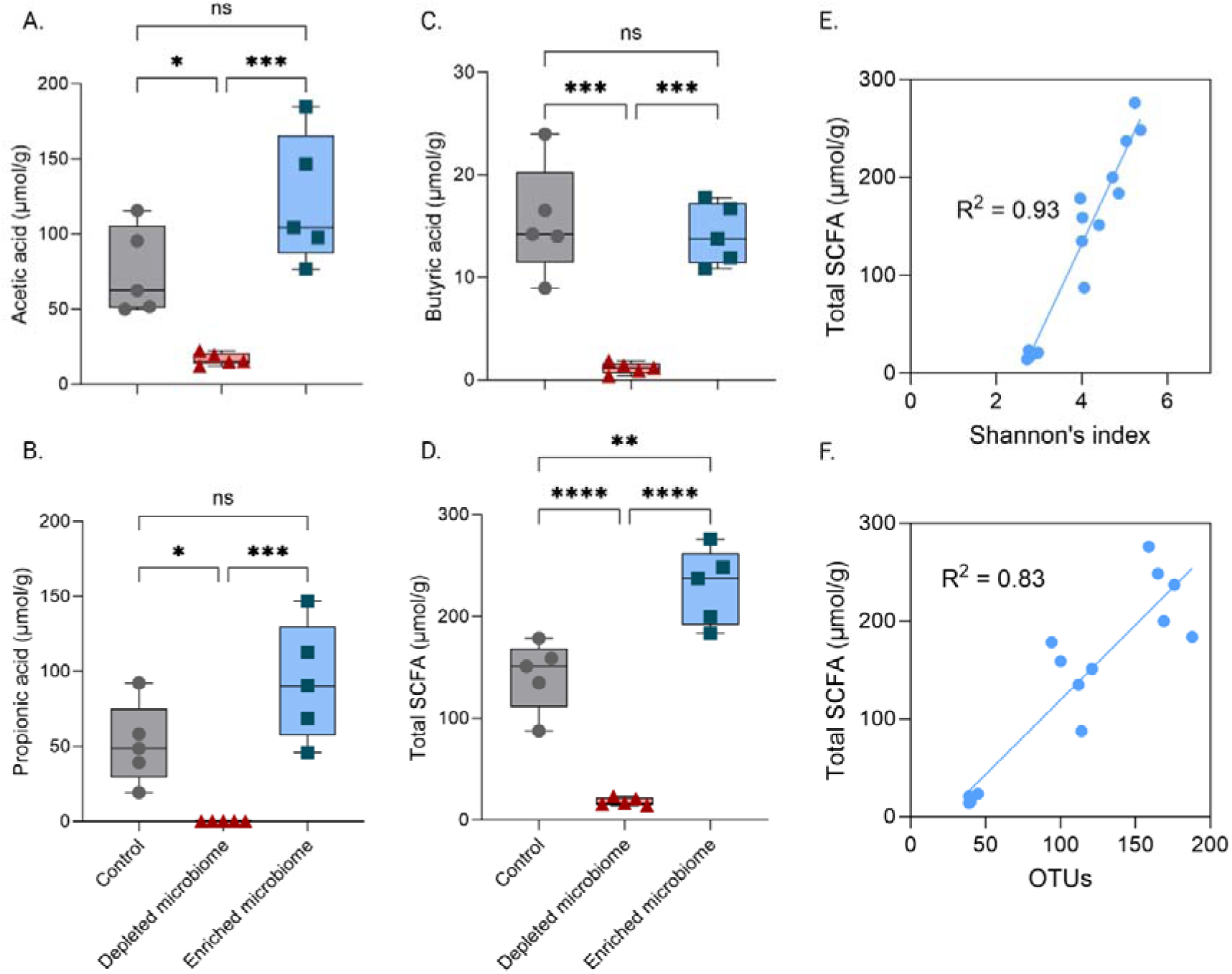
Enriched microbiota diversity correlates directly with an increase in SCFA concentrations in rats. [**A**] Acetic acid concentration across control, depleted, and enriched microbiome groups. [**B**] Propionic acid concentration across the three groups. [**C**] Butyric acid concentration across the three groups. [**D**] Total SCFA concentration across the three groups, demonstrating statistical significance across all three groups investigated. Total SCFA concentration directly correlates with alpha diversity, as demonstrated through strong correlations with Shannon’s index, showing strong correlation (R^2^ = 0.93) (Panel [**E**]) and OTUs (R^2^ = 0.83) (Panel [**F**]). Values are presented as the mean ± standard deviation (SD). Statistical significance was assessed using an unpaired t-test and one-way ANOVA. The asterisk (*), (**), (***) or (****) denote statistically significant results (p < 0.05), (p< 0.01), (p < 0.001) or (p < 0.0001) respectively.

### 3.3 The gut microbiome mediates the oral bioavailability of lurasidone

The pharmacokinetic profiles for each group following a single oral dose of lurasidone (20 mg/kg) to male Sprague Dawley rats are displayed in **Fig. 3A** with interindividual pharmacokinetic variability highlighted in **Fig. 3B-D**. Noteworthy is the outlier present in the depleted microbiome group (**Fig. 3C**) which showcases uncharacteristically prolonged lurasidone absorption/ delayed clearance in contrast to the other four rats. When excluded from consideration, the mean lurasidone bioavailability is diminished compared to the control group. The enriched microbiome group demonstrates a 4.3-fold increase in lurasidone bioavailability over the 24 hours in comparison to the control, based on mean area-under-the-curve (AUC_0-24_) values, presented in **Fig. 3E**. Further, C_max_ values increased by 3.3-fold and 3.9-fold for the enriched microbiome group compared to the control and depleted microbiome group, respectively (**Fig. 3F**). The degree of drug absorption between the control and depleted microbiome groups were statistically equivalent, but significant differences were observed in T_max_ values. Pharmacokinetic parameters are summarised in Table 1.

**Figure 3.**
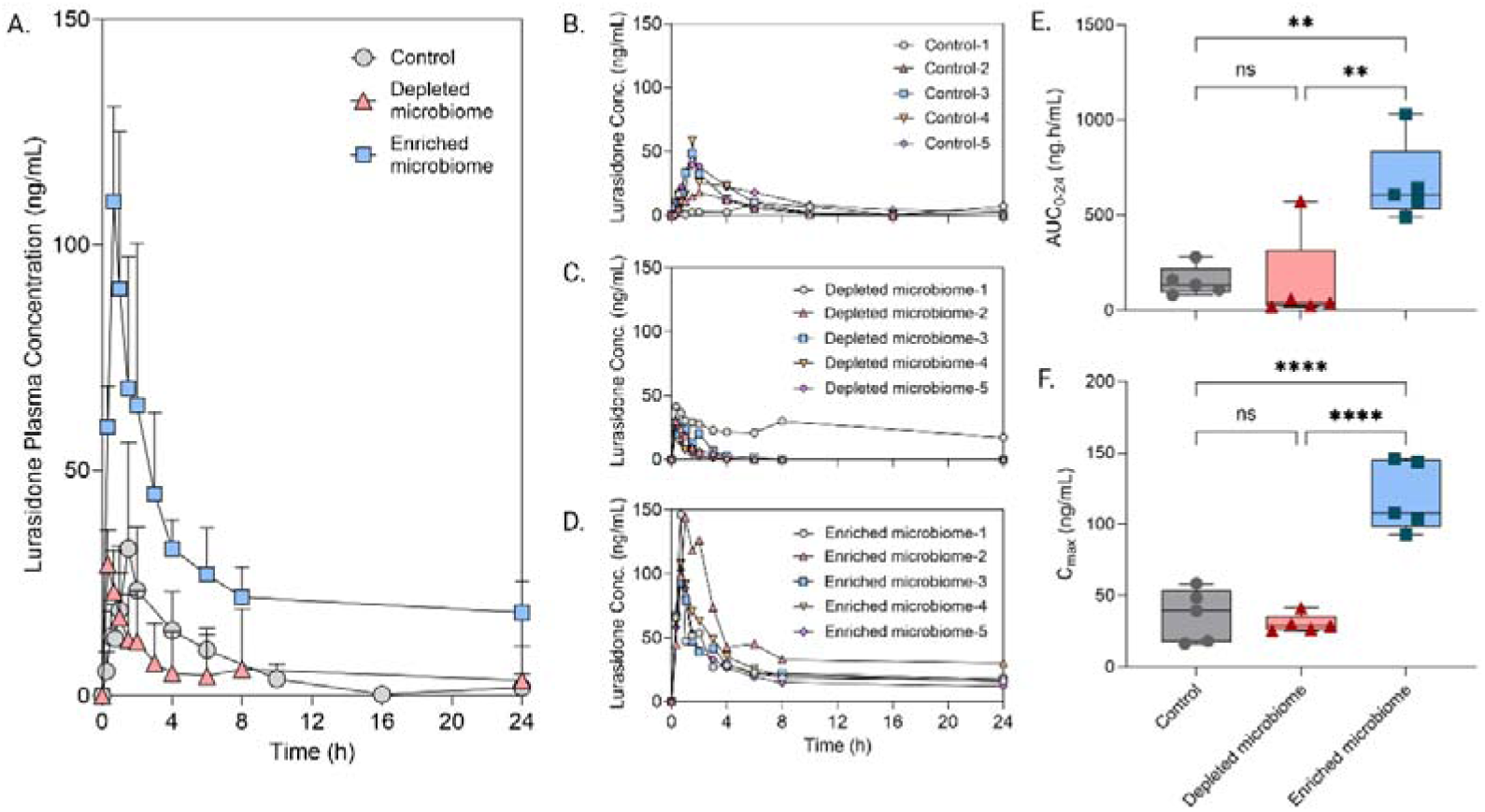
Microbiome enrichment via prebiotic supplementation increases lurasidone bioavailability. [**A**] Pharmacokinetic profiles for the oral administration of lurasidone (20 mg/kg) across the three groups (n = 5) revealed an increase in oral bioavailability for the enriched microbiome group. Panels [**B-D**] present the pharmacokinetic analysis of each rat within the three groups. [**E**] Showcasing area-under-the-curve across the groups with statistical significance observed between enriched microbiome and depleted microbiome groups, and enriched microbiome and control groups respectively. [**F**] Maximum lurasidone plasma concentration observed across the three groups trialled. Values are presented as the mean ± standard deviation (SD). Statistical significance was assessed using an unpaired t-test and one-way ANOVA. The asterisk (**) or (****) denote statistically significant results (p< 0.01) or (p < 0.0001) respectively.

**Table 1:**
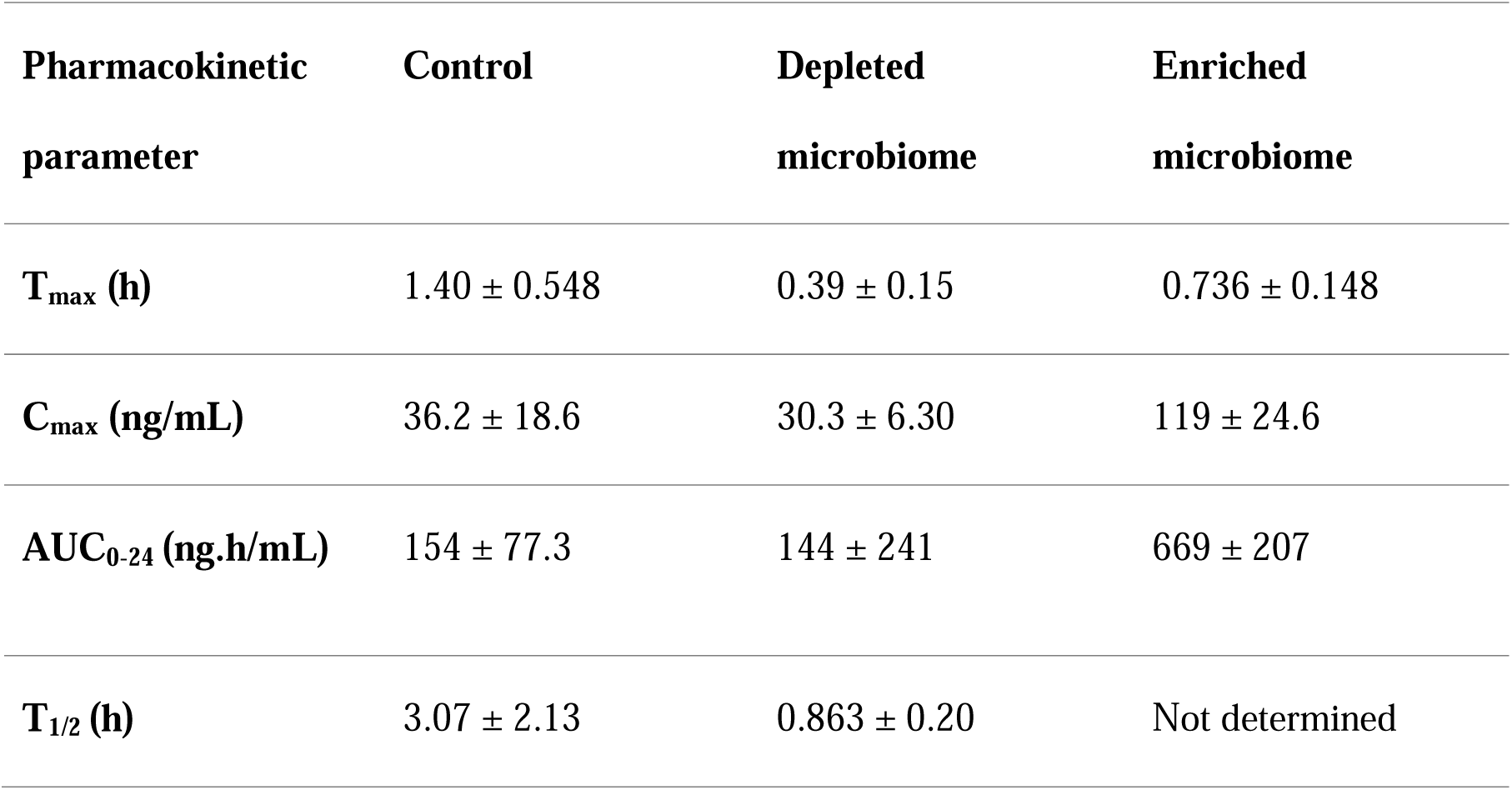
Pharmacokinetic parameters following oral administration of lurasidone (20 mg/kg) to male Sprague Dawley rats in the fasted state following a 14-day pre-treatment with antibiotics and prebiotics to deplete and enrich the gut microbiome, respectively. Values represent the mean ± SD, n = 5.

| Pharmacokinetic<br>parameter | Control | Depleted<br>microbiome | Enriched<br>microbiome |
| --- | --- | --- | --- |
| $T_{\max}$ (h) | $1.40 \pm 0.548$ | $0.39 \pm 0.15$ | $0.736 \pm 0.148$ |
| $C_{\max}$ (ng/mL) | $36.2 \pm 18.6$ | $30.3 \pm 6.30$ | $119 \pm 24.6$ |
| $AUC_{0-24}$ (ng.h/mL) | $154 \pm 77.3$ | $144 \pm 241$ | $669 \pm 207$ |
| $T_{1/2}$ (h) | $3.07 \pm 2.13$ | $0.863 \pm 0.20$ | Not determined |

### 3.4 Lurasidone bioavailability correlated with SCFA-mediated changes to intestinal pH by altering drug solubilisation

The impact of microbiome compositional changes on luminal pH are highlighted **Fig. 4A**. demonstrating significant alteration to intestinal pH accompanied due to gut microbiome manipulation. An enriched microbiome resulted in a lower luminal pH (mean pH ∼6.25), and a depleted microbiome revealed a higher luminal pH (mean pH ∼7.50), compared to the control group exhibiting a mean pH range of ∼6.75. Strong negative correlations existed between intestinal pH and Total SCFA (**Fig. 4B**) and Shannon’s Index (**Fig. 4C**), demonstrating that the intestinal pH decreases as a function of increasing SCFA concentration and microbiota diversity, respectively. Lurasidone solubility was investigated at equivalent pH ranges to those observed in *ex vivo* intestinal media, by incubating lurasidone within FaSSIF media at concentrations in excess of its solubility. **Fig. 4D** demonstrates a ∼2-fold increase in lurasidone solubility within FaSSIF media simulating an enriched microbiome (*i.e.*, at pH 6.25), compared to the control pH of 6.75. Lurasidone solubility at pH 7.5 is significantly diminished with an average equilibrium solubility of 2.22 ± 0.09 µg/mL. **Fig. 4E** and **4F** demonstrate strong positive correlations between area-under-the-curve and total SCFA (R^2^ of 0.72) and Shannon’s index (R^2^ of 0.71), respectively, highlighting the link between lurasidone bioavailability and metabolome and microbiota composition.

**Figure 4.**
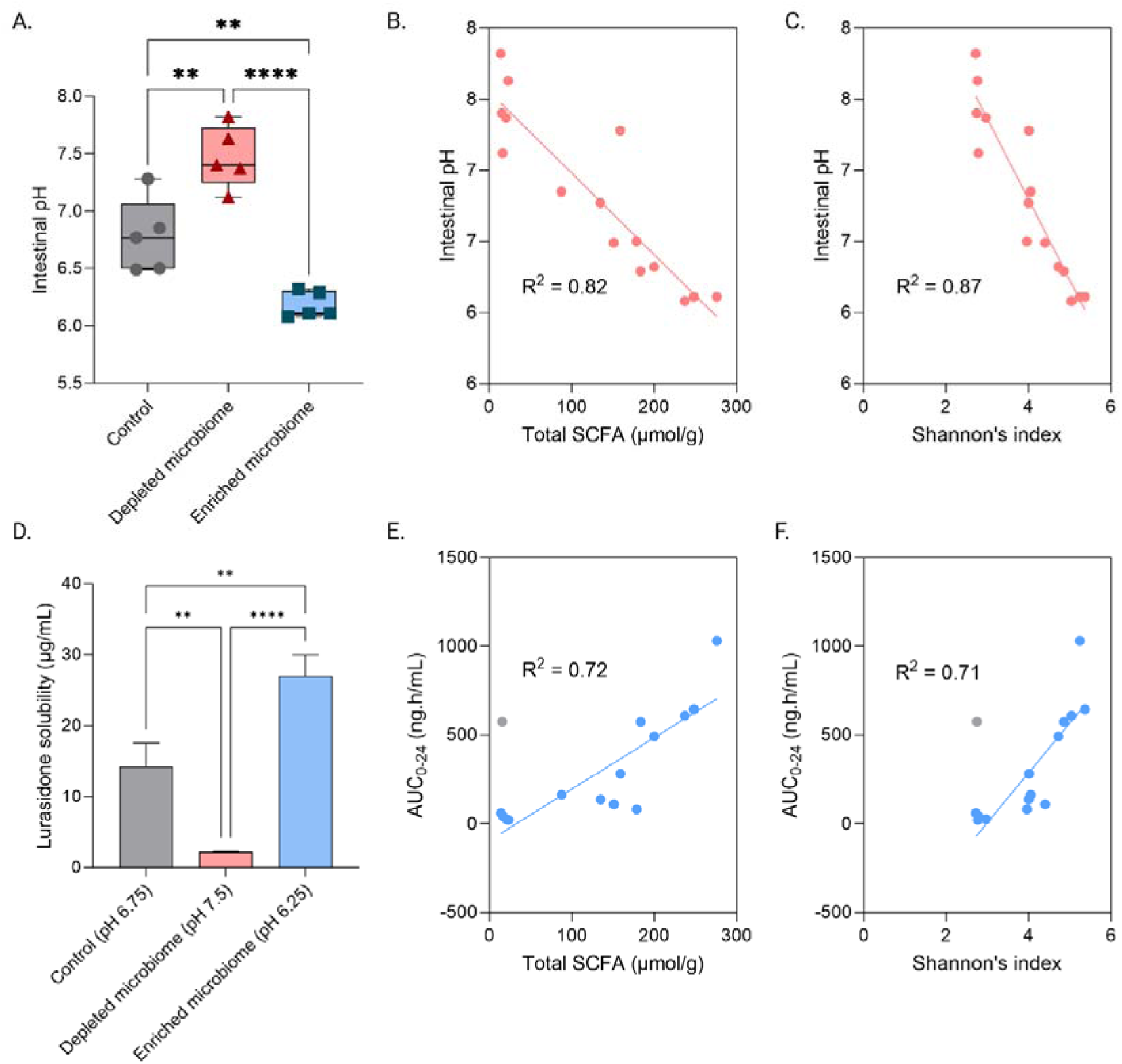
An enriched microbiome demonstrates increased lurasidone solubility due to a lower intestinal pH, facilitating increased lurasidone solubilisation. [**A**] Intestinal pH levels across control, depleted microbiome, and enriched microbiome, with the enriched microbiome displaying a statistically significant reduction in luminal pH. [**B** and **C**] A strong relationship exists between intestinal pH and both total SCFA concentrations (R^2^ = 0.82) and Shannon’s Index (R^2^ = 0.87). [**D**] Lurasidone solubility varies significantly in simulating intestinal media mimicking microbiome-induced pH changes, with prebiotic pretreatment leading to an enriched microbiome significantly increasing lurasidone bioavailability. [**E**] and [**F**] A strong correlation exists between lurasidone area-under-the-curve and both total SCFA (R^2^ = 0.71) and Shannon’s index (R^2^ = 0.72) Values are presented as the mean ± standard deviation (SD). Statistical significance was assessed using an unpaired t-test and one-way ANOVA. The asterisk (**) or (****) denote statistically significant results (p< 0.01) or (p < 0.0001) respectively.

## 4. Discussion

Pharmacomicrobiomics, the study of bidirectional relationships between drugs and the gut microbiome, is particularly relevant when prescribing antipsychotic medication as it underscores much of the observed inter-, intra-individual variability in pharmacokinetics and side effect profiles. [1,11,12,24]. While the impact of antipsychotics on the gut microbiome is gaining attention, as evidenced by comprehensive reviews from Minichino et al., the reciprocal relationship (*i.e.* the impact of the gut microbiome on antipsychotic pharmacokinetics) is less explored [1,3]. Therefore, this study investigated the impact of the gut microbiome on lurasidone pharmacokinetics using an *in vivo* rat model where animals’ gastrointestinal microenvironments were manipulated through pre-exposure to prebiotics, antibiotics, or a control group (as per **Fig. 1A**).

### 4.1 Gut Microbiome Alteration and Bacterial Composition

Antibiotic supplementation decreased several commensal bacterial families, namely *Lachnospiraceae* and *Bacteroidaceae* as evidenced by **Fig. 1B-C**. Similarly, antibiotic supplementation significantly depleted the gut microbiome, leading to a halving of OTUs (**Fig. 1D)** and likewise halving of microbial α-diversity (represented by Shannon’s index: **Fig. 1E**). Such depletion of the gut microbiome illustrates a stark decrease in ecosystem capacity and is consistent with previous observations by Langdon et al. [25]. Conversely, enrichment of the gut microbiome increased both *Lachnospiraceae* and *Bacteroidacae* relative abundance (**Fig. 1B-C**.) alongside holistic improvements in OTUs (**Fig. 1D**) and increases in β-diversity (**Fig. 1F**) both of which align with previous studies by Carlson et al. and Tian et al. [26,27]. Findings from Meola et al. suggest the enrichment protocol used (inulin at 100 mg/L) allowed for commensal microbial fermentation and thus promoted a eubiotic state in the gut ecosystem [16].

### 4.2 Relationship between Gut Microbiome and Short-Chain Fatty Acid Production

SCFAs (acetate, propionate, and butyrate) are primarily produced through gut bacterial fermentation of dietary fibres in the colon and are vital for the maintenance of gut integrity, immune modulation and gut-brain axis signalling amongst other roles [18,28,29]. In line with current literature, increases in overall OTUs led to increases in SCFA concentrations (**Fig. 2D**) with strong correlations being noted between SCFA concentration and Shannon’s index and OTUs (**Fig. 2E**: R^2^ 0.93, **Fig. 2F**: R^2^ 0.83). Such results are echoed by Gurry et al. with similar correlations between functional heterogeneity in gut microbiome and SCFA production [30].

### 4.3 Impact of Short-Chain Fatty Acids on Luminal pH

The relationship between intestinal pH and SCFA concentration (explored in **Fig. 4B**) demonstrated a strong correlation (R^2^: 0.82) whereby an increase in SCFA concentration led to a decrease in intestinal pH and is consistent with findings by Walker et al. [17]. Likewise, gut microbial heterogeneity also correlates with intestinal pH where enriched and depleted microbiomes (**Fig. 4A**) showcased contrasting results (enriched microbiome pH: 6, depleted microbiome pH: 7.5). Liu et al. explored mechanistic reasoning for this observation, suggesting the acidic nature of SCFAs (propionate: pH 2.4, pKa 4.87, butyrate: pH 5.5 pKa 4.82, acetate: pH 4.6, pKa 4.76) can decrease luminal pH through the release of H□ ions [31]. Specifically, the acidity of SCFAs arises from the presence of a carboxyl group (-COOH), which can donate a proton (H□) to the solution, forming a carboxylate anion (R-COO□) stabilised by resonance as suggested by Parada Venegas et al. [18]. The inductive effect of the alkyl chains (methyl, ethyl and propyl) also slightly influences their acidity, with longer chains being less acidic due to electron-donating impacts [18]. Their acidic nature is explored in a physiological context by Sun et al. suggesting this luminal pH reduction to be integral in inhibiting pathogenic invasion and colonisation [29].

### 4.4 Impact of Luminal pH on Lurasidone Solubility and Bioavailability

In the context of pharmacomicrobiomics, the acidic environment can drive drugs toward or further from their pKa, influencing drug charge and thus solubility within the GI tract, as noted by Panebianco et al. [24]. Lurasidone demonstrated variable pH-dependent solubility during *in vitro* studies that simulated the intestinal fluid of microbiome depleted and enriched animals (**Fig. 4D**). The lower pH environment in the presence of an enriched microbiome increased lurasidone solubility to 30 µg/mL (compared to 15 µg/mL and 3 µg/mL for the control and depleted microbiomes, respectively). The current findings correlate well with a previous study by Qian et al. that has highlighted the pH-dependent solubility of lurasidone in simulated intestinal fluids [14]. Lurasidone contains a benzothiazole derivative with a piperazine ring (providing basicity and contributing to pKa), a sulphonamide group (neutral but capable of influencing hydrogen bonding capacity), and multiple aromatic rings (impacting lipophilicity and drug-receptor interactions), resulting in a pKa of 7.6 [13]. Below the pKa, the piperazine nitrogen is protonated, resulting in a positively charged species (Lurasidone-NH□□) which possesses greater aqueous solubility and hydrophilicity, through the formation of ion-dipole interactions between the positively charged amine and water molecules [32,33]. This results in the maximum aqueous solubility of 0.349 mg/mL in pH 3.5 buffer [13]. In this context of the observations made, such protonation may underscore the increased solubilities at lowered pH observed in enriched microbiomes (**Fig. 3A**).

Ultimately, the 2- to 10-fold increase in lurasidone solubility for the microbiome enriched group cumulated in a >4.3-fold increase in lurasidone bioavailability (illustrated in **Fig. 3A**), based on AUC values (**Table 1**). On average, depleting the gut microbiome through antibiotic pre-treatment did not alter the oral bioavailability compared to the control group (**Fig. 3A**). However, it is important to note that when comparing the pharmacokinetic profiles of individual animals (**Fig. 3B-D**), it is evident that ‘Depleted microbiome profile-1’ represents an outlier with greater than expected lurasidone-plasma concentrations. When taking this into account, lurasidone bioavailability is significantly reduced through antibiotic pre-treatment with strong correlations evident between SCFA concentration and AUC_0-24_ for lurasidone (depicted in **Fig. 4E**). Thus, based on these findings, it can be concluded that the microbiome-modulation of SCFA concentrations, and subsequent changes in intestinal pH, play an integral role in solubilising the PWSB lurasidone, enabling its absorption into the systemic circulation, as schematically depicted in **Fig 5**.

**Figure 5.**
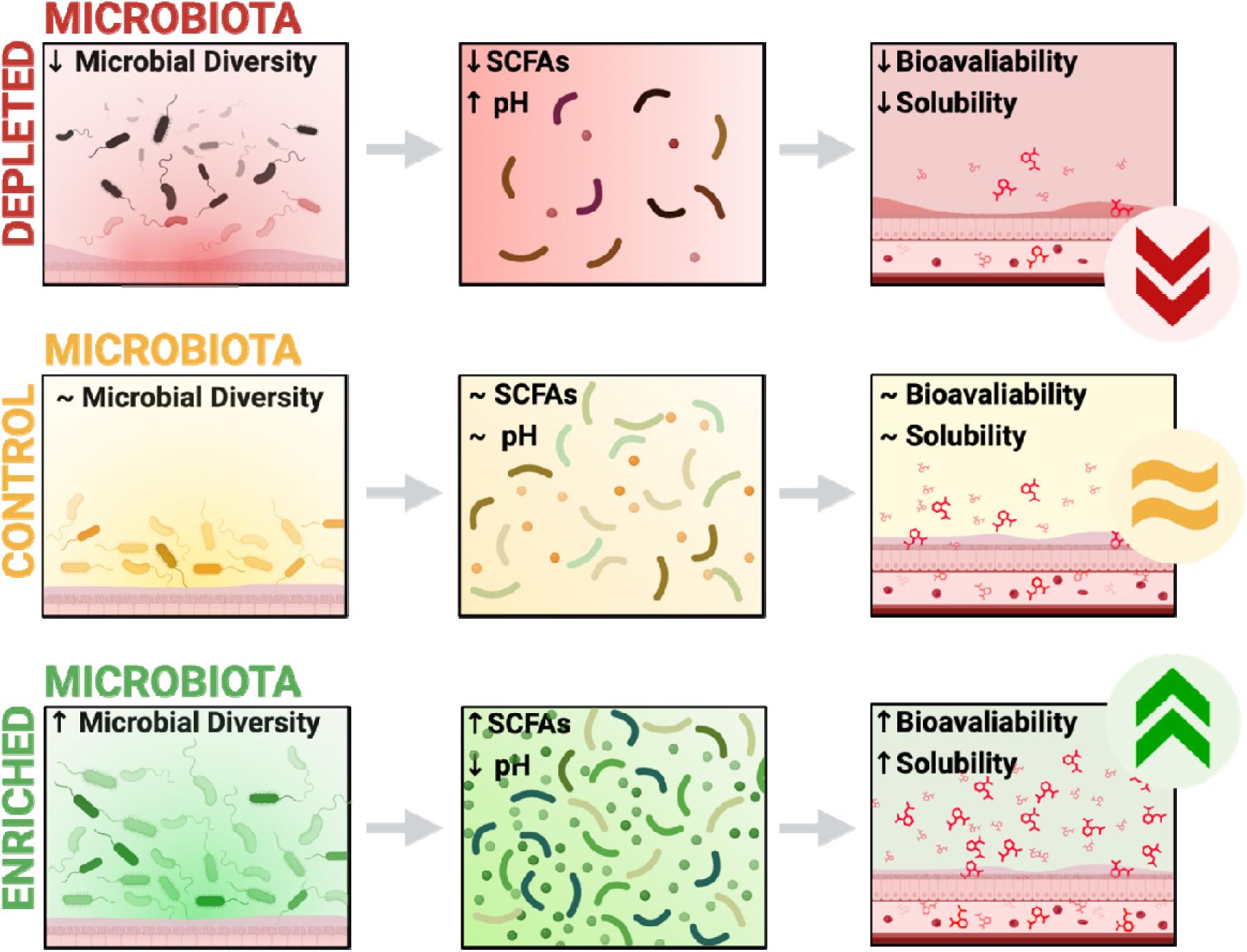
Gut microbiome manipulation with antibiotics and prebiotics alters intestinal pH through modulation of short-chain fatty acid abundance. These changes to the gastrointestinal environment drive changes to the solubility and thus absorption of the poorly soluble weak base, lurasidone, whereby an enriched microbiome leads to greatest oral bioavailability.

Whilst the specifics of lurasidone absorption and active transporter receptors are poorly understood, increased solubility is generally accepted as being favourable towards increasing absorption and bioavailability (as corroborated by **Fig. 3E**) and is thoroughly explored in a review by Kumari et al. [34]. The main mechanism suggested is the increased concentration gradient across the intestinal epithelium and the potential for paracellular absorption of the charged species [34]. Whilst not previously explored for lurasidone, similar relationships are summarised in a comprehensive review by Minichino et al. [3].

Alternative atypical antipsychotics also possess similar pH-dependent solubility characteristics, such as olanzapine (protonation of amine group) and risperidone (protonation of the piperidine nitrogen) [35]. However, unlike alternatives, lurasidone is hampered by poor bioavailability (9-19%) with a 2-fold positive food effect owing to its poor aqueous solubility [9]. Such findings may explain why lurasidone prescription rates are considerably lower than alternatives such as olanzapine (*e.g.,* 111,911 prescriptions for lurasidone in Australia compared to 719,915 for olanzapine) [36].

### 4.5 Implications and Future Directions

The novel findings suggest a new potential contributor to lurasidone inter- and intra-patient pharmacokinetic variability, namely the gut microbiota and metabolome. These findings are corroborated by emerging work from Zhao et al. and Tsunoda et al. who have demonstrated the gut microbiome composition to be capable of predicting drug pharmacokinetics [37,38]. Ultimately, the correlations between SCFA, intestinal pH and subsequent solubilisation and absorption of lurasidone observed, illustrate a future wherein predictive models may use SCFAs as accurate biomarkers for managing personalised therapy. Currently, SCFAs are tested as putative indicators for colorectal and non-small cell lung cancers [39]. However, these scenarios are limited in scope, restricted to merely suggesting gut inflammation characteristics and immune activation [40].

While the current work demonstrated correlations between gut microbiome composition, SCFA abundance, intestinal pH and lurasidone bioavailability, several other pharmacokinetic mechanisms exist that can dictate changes in drug-plasma concentrations. For example, the current study ignores the role of drug metabolism on bioavailability. Previous studies have demonstrated the capacity for the gut microbiome to modulate hepatic cytochrome P450 (CYP450) enzyme activities [41,42]. In the context of lurasidone, this presents another avenue of pharmacokinetic modulation as lurasidone is extensively metabolised (through oxidative N-dealkylation between the piperazine and cyclohexane rings) by CYP450, forming lurasidone metabolites 14326, 14283 (both active) and 20219, 20220 (inactive) [43]. Thus, future studies should focus on understanding the impact of gut microbiome modulation on CYP450 activity in the context of lurasidone metabolism, to provide a more complete picture of the mechanisms driving changes in lurasidone absorption, beyond changes to the metabolome.

Given the role of intestinal pH and the gut microbiome in modulating lurasidone solubility and absorption, it may necessitate a re-evaluation of treatment regimens for schizophrenia to accommodate gut microbiome interventions (such as enrichment using prebiotics) to maximise pharmacotherapy efficacy. For example, prebiotic co-therapy has already been explored for impacts on drug characteristics (as explored by Kao et al. to mitigate olanzapine-induced weight gain), however the observations made in this study suggest both an underlying mechanism and scope for exploring new approaches for mitigating pharmacokinetic variability and thus, improving patient response rates to antipsychotics [44]. Further, such investigation may revitalise lurasidone therapy, which is made pertinent by recent evidence that lurasidone presents ‘gut neutral’ characteristics and imparts generally favourable impacts on the gut microbiome, thus it may serve as a better-suited therapeutic option for metabolic sensitive patients [5].

Furthermore, it is hoped that the current findings provoke formulation and delivery strategies to maximise lurasidone solubility, absorption, and bioavailability. An innovative example lies in the work by Meola et al. through the formulation of a lipid-based nanostructure and porous silica nanostructure which enhanced bioavailability of lurasidone by 8.8-fold [16]. Meola et al. also trialled inulin-lipid microcapsules to target the gut microbiome and improve lurasidone plasma concentration by 3+ fold whilst also enriching the microbial composition. Such optimised therapy led to 2.2-fold increases in systemic serotonin concentrations compared to pure drug which coincided with a 1.6-fold increase in faecal serotonin levels [16]. Similarly, Gamboa-Arancibia et al. formed a lurasidone-hydroxypropyl-β-cyclodextrin (obtained through the hydroxylation of β-cyclodextrin) and observed increases in solubility (up to 20 mg/mL) attributable to the presence of the hydroxyl substituent in this cyclodextrin [10]. Cyclodextrins can encapsulate hydrophobic drugs within their lipophilic cavity, while their hydrophilic exterior improves aqueous solubility. Thus, based on these emerging formulations and the results observed, it is expected that the current findings will guide future research in optimising therapy and potentially lead to increased prescription and clinical use of lurasidone in standard therapy for treating schizophrenia and bipolar disorder.

## 5. Conclusion and Prospects

For the first time, the intricate relationships between the gut microbiome, short-chain fatty acid (SCFA) production and luminal pH have been shown to exert a strong influence on the solubility and bioavailability of a poorly soluble weak base drug, lurasidone. Changes in microbiome composition, including key families of bacteria such as *Lachnospiraceae* and *Bacteroidaceae*, alters the intestinal environment by influencing SCFA production, which consequently affects luminal pH. This potentiates the observed inter- and intra-patient variability in lurasidone pharmacokinetics. Such inconsistencies may come down to nuances in gut microbiome compositions which ultimately influence luminal pH and drug solubility. As such, the novel observations made suggest the gut metabolome, namely SCFA concentrations, to be a key biomarker relevant for predictive modelling of lurasidone response.

Given lurasidone is ‘gut neutral’ and triggers minimal metabolic dysfunction, it serves as a better-suited therapeutic option for metabolically sensitive patients. Thus, promoting its place in therapy is imperative as is exploring strategies to optimise drug delivery. Recent successes with lipid-based nanostructures and cyclodextrin complexes in improving lurasidone’s solubility are promising, potentially addressing bioavailability challenges and leading to more effective treatments for schizophrenia and bipolar disorder. Furthermore, the implications extend beyond lurasidone to other pH-sensitive drugs, suggesting advancements in drug development and administration across various therapeutic areas. By considering the gut microbiome’s role in drug pharmacokinetics, pharmaceutical research may uncover new strategies to optimise drug delivery and efficacy.

The critical role of the gut microbiome in antipsychotic efficacy highlights new avenues for improving mental health treatment. Bridging microbiology, pharmacology, and psychiatry paves the way for innovative, microbiome-informed approaches, ultimately improving patient health outcomes through the promotion of alternatives such as lurasidone.

## Acknowledgements

The Hospital Research Foundation (THRF) Group is gratefully acknowledged for their EMCR Fellowship funding and generous support provided to Dr Paul Joyce (2022-CF-EMCR-004-25314). The authors acknowledge the University of Adelaide, enabled by NCRIS, university, and state government support.

